# Structured connectivity exploits NMDA non-linearities to enable flexible encoding of multiple memoranda in a PFC circuit model

**DOI:** 10.1101/733519

**Authors:** Stefanou S. Stefanos, Athanasia Papoutsi, Panayiota Poirazi

**Affiliations:** Institute of Molecular Biology and Biotechnology (IMBB), Foundation for Research and Technology-Hellas (FORTH), Heraklion, Crete, 70013, Greece; Department of Biology, University of Crete, Heraklion, Crete, 70013, Greece

**Keywords:** prefrontal, working memory, dendritic integration, NMDA, behavioral flexibility, assemblies, dimensionality

## Abstract

Prefrontal Cortex (PFC) exerts control on action selection and mediates behavioral flexibility during working memory (WM) task execution, when integration and retention of sensory information takes place. We used biophysical circuit modelling to investigate the dendritic, neuronal and circuit mechanisms that underlie these computations, aiming to causally link these three processing levels. Our simulations predict that dendritic NMDA non-linearities drive distinct activity dynamics of the same network, thus enabling adaptive coding in the absence of plasticity mechanisms. Specifically, we find that distinct assemblies of mixed-selectivity neurons emerge and fire in stable trajectories in a stimulus-dependent manner. Synaptic inputs that are spatio-temporally clustered, as provided by the structured connectivity of the PFC, facilitate these activity dynamics, thus further increasing the flexibility of the network. Our study suggests that behavioral flexibility may result from the formation of memoranda-specific assemblies in the PFC which are utilized dynamically in relation to the task at hand.

## Introduction

Working memory (WM) is the ability to maintain behaviorally relevant stimuli for a short period of time and to integrate them towards a choice/action. Seminal WM studies found that the activity of single neurons increases during the presentation of a to-be-maintained item (stimulus epoch) and persists throughout the blank ‘post-stimulus’ interval (post-stimulus epoch). This persistent activity constitutes a basic component of decision making by bridging the temporal gap between the item presentation and the subsequent contingent response (Funahashi et al., 1989; Fuster and Alexander, 1971) and is involved in the top-down modulation of behavior (Van Kerkoerle et al., 2017). The above findings form the backbone of the so-called fixed selectivity framework that has dominated the WM field. According to this framework, selective tuning of neurons or neuronal populations to the to-be-remembered information can be explained by the formulation of stable attractors: the fine-tuning of the respective network connections creates stable points, whereby individual neurons fire consistently to specific stimuli, both during the stimulus and the post-stimulus epoch (Renart et al., 2003; Wimmer et al., 2014).

Recent electrophysiological findings challenge this framework, based on the observation that the dynamics underlying WM in the PFC are more diverse than previously thought. First, individual neurons display time-varying discharge rates (Bernacchia et al., 2011; Brody et al., 2003; Jun et al., 2010; Shafi et al., 2007), leading to the hypothesis that the tuning properties of neurons change throughout the task duration. Second, individual PFC neurons can be selective for a broad range of task variables and stimuli, exhibiting thus a ‘mixed selectivity’ (Lindsay et al., 2017; Rigotti et al., 2013; Warden and Miller, 2010). Third, at the population level, the stimulus-induced network activity decays after the stimulus offset and emerges later with a different population activity pattern (Stokes et al., 2013), yet containing a stable representation of the to-be-remembered item (Murray et al., 2017). These findings challenge the fixed-selectivity framework and call for a deeper mechanistic understanding of the cellular substrate of WM.

Single-neuron dynamic behavior and mixed selectivity has thus far been implemented using recurrent neural networks with random connectivity and non-biologically plausible training of either the weights of the readout neuron or of the entire network (Barak et al., 2013a, 2013b; Mante et al., 2013; Rigotti et al., 2010). Unlike the slow training required by these approaches and the resulting rigidity in responses, PFC circuits typically utilize short-term dynamic mechanisms that renders them highly flexible (Bouchacourt and Buschman, 2019; Fujisawa et al., 2008). In addition, the representational shift at the population level from the sensory towards a memory state has thus far been attributed to either the dynamic behavior of a randomly connected recurrent neuron network (Bouchacourt and Buschman, 2019) or to short-term facilitation mechanisms (Barak et al., 2010). Both approaches depend on the implementation of distinct populations of sensory and memory neurons. Yet, experiments show that selection and integration are implemented by the same PFC circuit, with both representations being present at the single neuron level (Mante et al., 2013; Warden and Miller, 2010). In this case, the shift in the population activity indicates integration of an incoming stimulus with the network’s internal state, which depends on short-term dynamics (Buonomano and Maass, 2009).

At these short-term time scales, it remains an open question how a recurrently connected network utilizes a rapid mechanism that renders it able a) to perform both selection and integration b) to be highly dynamic at the single neuron level yet, c) stable and separable at the population level. Theoretical work suggests that the answer may lie in the highly non-random connectivity of pyramidal neurons, characterized by distance-dependent formation of neuronal clusters and over-represented structural ‘motifs’ (structured connectivity) (Perin et al., 2011; Song et al., 2005), as this type of connectivity is optimal for storing a large number of attractor states in a robust fashion (Brunel, 2016). In the PFC in particular, layer 5 (L5) pyramidal neurons have almost double the reciprocal connections of other cortical regions, forming hyper-reciprocally connected circuits (Morishima and Kawaguchi, 2006; Morishima et al., 2011; Wang et al., 2006). This hyper-reciprocity could be a major player in the formation of multiple, transient, assemblies in the PFC. In addition to connectivity features, the NMDA receptor is considered a key determinant of stable attractor dynamics (Durstewitz et al., 2000; Wang, 1999). The non-linearity and slow kinetics of the NMDA receptor make it a complementary candidate to the short-term plasticity mechanisms, that can contribute to the activity-dependent selectivity and storage of the sensory input i.e. adaptive coding (Stokes et al., 2013). In support of this hypothesis, NMDA-dependent dendritic non-linearities have been shown to underlie the expression of memory states in computational models of small cell assemblies (Papoutsi et al., 2014, 2017).

In this work we investigated the role of NMDA-dependent dendritic nonlinearities and structured connectivity in the ability of a biophysical PFC network model to express multiple distinct memoranda. Towards this goal, we use a detailed biophysical network model of the PFC that is heavily constrained in terms of its connectivity and biophysical properties and show that it reproduces multiple types of experimental data. Overcoming many of the shortcomings of previous implementations, this model incorporates several recent findings of PFC function. Specifically, our model responds with single neuron variability comparable to what is experimentally reported (Bernacchia et al., 2011; Brody et al., 2003; Jun et al., 2010; Shafi et al., 2007). Moreover, at the population level, the model exhibits the representation shift between the stimulus and memory epochs, while maintaining robust memoranda over time (Murray et al., 2017; Stokes et al., 2013). This is achieved without the need to artificially simulate different sensory and memory populations, rather utilizing a recurrent neural network demonstrating that stimulus selection and integration can occur in a single network (Mante et al., 2013; Warden and Miller, 2010). We predict that NMDA non-linearities serve as a complementary mechanism for adaptive coding in the PFC, supporting diverse activities of a single population of the PFC, ultimately to be utilized for different behaviors.

## Results

### A novel, biologically-constrained network model of the PFC

To investigate the cellular and connectivity mechanisms underlying the experimentally reported properties of WM, we developed a novel, biologically-constrained compartmental network model of the PFC. The network consisted of 250 biophysically detailed and morphologically simplified L5 PFC pyramidal neurons (PC) and 83 fast spiking interneurons (FS). Single neuron models were validated against experimental data with respect to both active and passive electrophysiological properties (Table S1). Basal dendrites of pyramidal neurons supported NMDA-dependent non-linear integration, as found experimentally (Figure 1A, Table S2). Non-random connectivity was implemented in congruence with available experimental data (Figure S1, Tables S3, 4). Specifically, the PC pair connection probability was set to be (a) distance-dependent and (b) local clustering dependent, based on the common neighbor rule (Perin et al., 2011; Song et al., 2005; Wang et al., 2006). PC-PC connections were created to reflect the high reciprocity ratio (47% of total PC-to-PC connections, given that there is a connection) found specifically in the PFC (Morishima et al., 2011; Wang et al., 2006) (Figure S1B-E). FS interneurons were connected to PCs in a sparse and distance-dependent manner (Packer and Yuste, 2011), respecting the connectivity reciprocity ratio for FS to pyramidal projections in L5 of the frontal cortex (Otsuka and Kawaguchi, 2009) (Figure S1F-H). In addition, 66% of the FS interneurons were connected with each other as reported in (Galarreta and Hestrin, 1999). Concerning network creation, see Methods, sections “Network Connectivity” and “Synaptic connectivity properties”.

**Figure 1.**
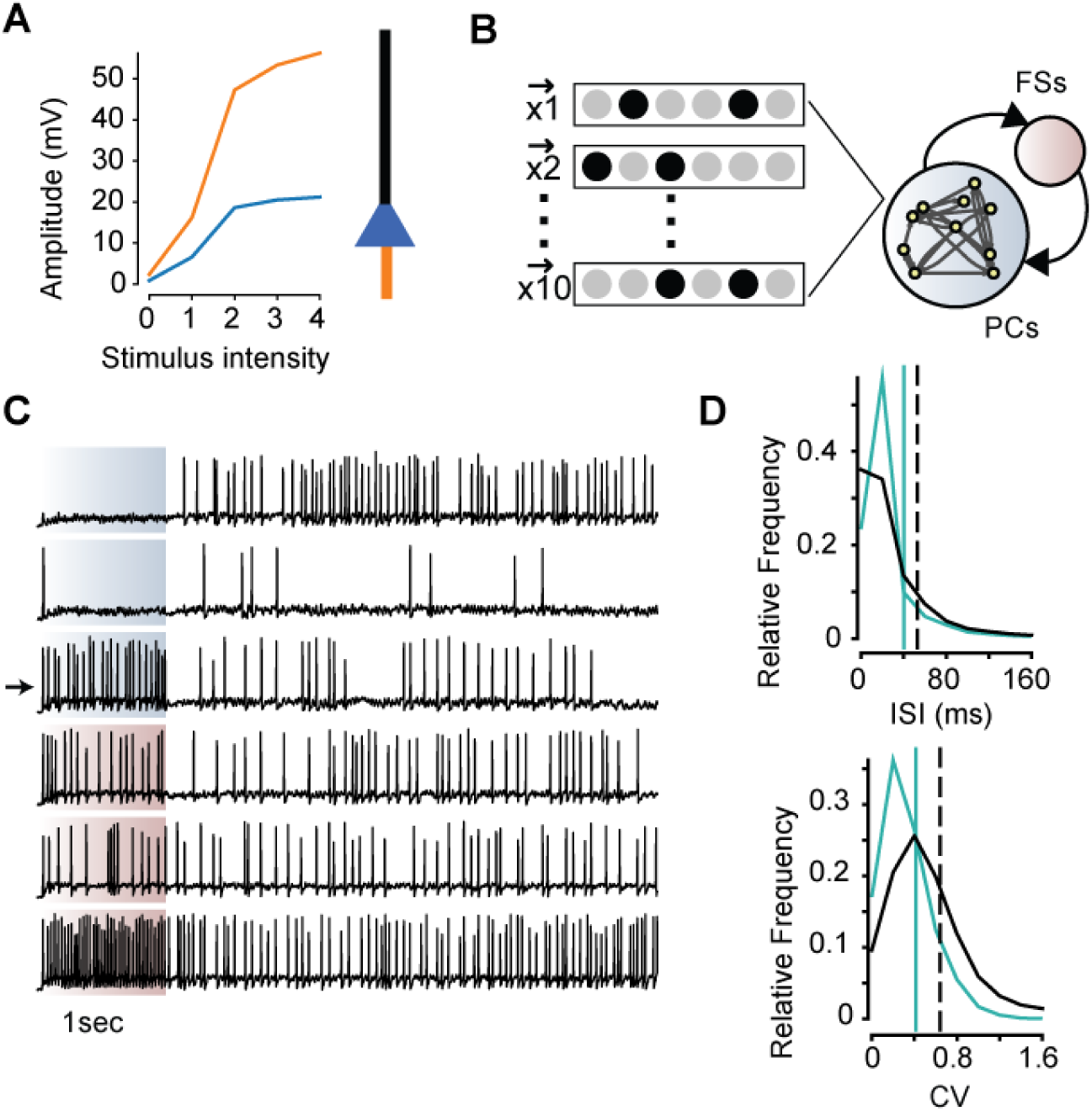
PFC network model reproduces single neuron dynamics. (A) Left: Non-linear NMDA responses are generated in the basal dendrites of the pyramidal neurons as in (Nevian et al., 2007). Orange: voltage response of the basal dendrite while increasing the number of activated synapses (2 events at 50Hz, activation of 1 to 21 synapses, with step 5). Blue: Corresponding somatic response. Right: Cartoon morphology of the pyramidal neuron model. Orange: basal dendritic compartment, blue: somatic compartment, black: apical dendritic compartments (B) Schematic representation of the network model. Fifty randomly selected PCs were stimulated in each trial, as indicatively denoted by the vectors **x**. (C) Exemplar voltage responses of PCs (top three) and FSs (bottom three) in a single trial. The shaded region indicates stimulus presentation (1sec). Arrow: PC that received stimulation. (D) Histograms of inter-spike intervals (ISIs) (top) and Coefficient of Variation of ISIs (CV, bottom) for PCs. Data are pooled for different structured network instances and trials for the stimulus period (green) and post-stimulus period (black). Distribution means are indicated with vertical lines.

### The model reproduces single neuron and population dynamics

To simulate WM in the network model, synaptic stimulation of 1-sec long, uncorrelated, Poisson-like spike trains was delivered to random subpopulations consisting of 50 PCs (20% of total, n=10 trials, Figure 1B). We focused on the emergence and dynamics of activity following the end of stimulus presentation (hereby termed post-stimulus epoch). Since both stimulated and non-stimulated neurons were recruited during the post-stimulus epoch with diverse spiking profiles (Figure 1C), we considered in our analysis all PCs and not just the stimulated ones. Trials with no PC activity during the last second of the simulation were not included in the analysis. Both inter-spike intervals (ISIs) and Coefficient of Variation (CV) were increased during the post-stimulus epoch (ISI_stimulus_=40.3±53.8ms, ISI_post-stimulus_= 52.9±115.3, p value=0.0004, Mann Whitney U test, CV_stimulus_=0.42±0.25, CV_post-stimulus_= 0.64±0.46, p value=0.0001, Mann Whitney U test, Figure 1D, ‘control’ cases), in line with experimental data (Compte et al., 2003; Stokes et al., 2013).

At the network level, activity was sparse (sparsity=0.89, values close to 1 correspond to sparser activity, see Methods, section “Sparsity”, (Rolls, 1997)) and characterized by dynamic population responses; the firing of some neurons persisted, increased or fluctuated during the post-stimulus period (Figure 2A) (Bernacchia et al., 2011; Brody et al., 2003; Jun et al., 2010; Shafi et al., 2007). To assess whether the model reproduces the experimentally reported shift from sensory to the memory representation(s) (Spaak et al., 2017), we measured the correlation coefficient of the binned network activity (bin size q=50ms) between different simulation time points. Figure 2B depicts the reduction in the correlation of the network activity during the stimulus and the post-stimulus epochs, denoting the representation shift. Within the stimulus epoch, population activity is consistently highly correlated, yet, during the post-stimulus epoch correlation is stronger over the diagonal time axis, indicating that population activity is not static. The population velocity (i.e. the speed with which the network activity changes, see Methods, section “Population multidimensional velocity”) drops upon stimulus withdrawal and remains at stable values during the post-stimulus period (Figure 2C, top). Note that changes in the energy state (i.e. energy velocity, change in the average firing frequency of the network, see Methods, section “Population multidimensional velocity”) are lower: they peak upon stimulus withdrawal and reach almost zero values during the post-stimulus period (Figure 2C, bottom). This indicates that the shift is not due to changes in the average firing rates per se, as seen also experimentally (Stokes et al., 2013). The above simulations replicate 1) the irregular spiking activity of single neurons reported in (Bernacchia et al., 2011; Brody et al., 2003; Jun et al., 2010; Meyers et al., 2008; Shafi et al., 2007) 2) the de-correlation of the network activity in the late post-stimulus epoch from the stimulus epoch reported in (Barak et al., 2010; Murray et al., 2017; Stokes et al., 2013) and 3) the change in the population activity trajectories over time (Stokes et al., 2013). Thus, our model network reproduces the complexity of single neuron and population responses during the post-stimulus epoch.

**Figure 2.**
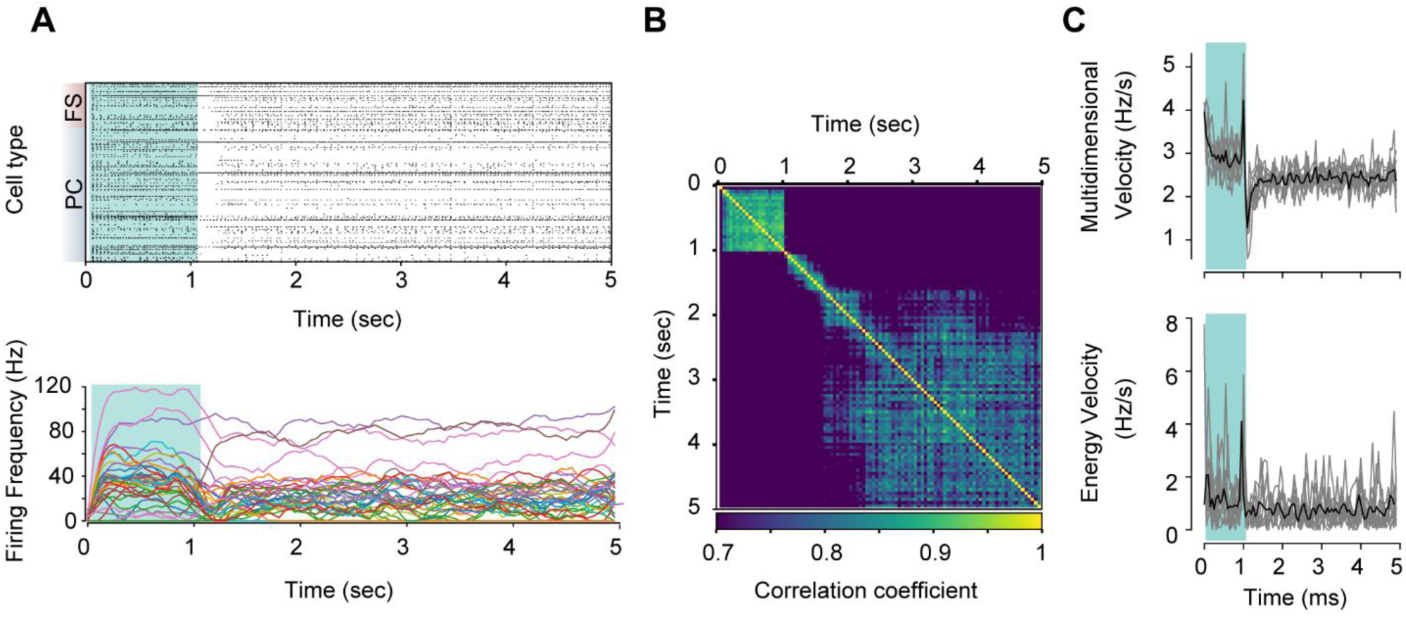
Dynamic population activity following stimulation. (A) Top: Raster plot of network activity in response to a 1-sec stimulus (shaded green area), for PCs and FSs. Bottom: Smoothed (temporal window: 11ms) firing frequencies of each PC, showing the different and highly dynamic responses over time (same trial as in A). Only PCs with average firing frequency across stimulus and post-stimulus epochs above 10Hz are shown here. (B) Cross correlation of network states (activity of PCs in a temporal window q=50ms) across the stimulus and post-stimulus epochs (aligned on stimulus onset, stimulus duration: 1sec). Color bar indicates the correlation coefficient, focused on the 0.7-1 range. (C) Top: Multidimensional velocity of network response, aligned on stimulus epoch onset. Bottom: Energy velocity (change in mean firing rate), aligned on stimulus epoch onset. Shaded region corresponds to stimulus epoch.

### The PFC network model expresses multiple memoranda

The shift from the stimulus-driven network activity to a different, memory-related one is more prominently observed in PCA state space (Figure 3A). Network responses for different trials, whereby for each trial a randomly selected sub-population of PCs was stimulated with uncorrelated Poisson inputs (see Methods, section “Stimulation”), diverge from the stimulus epoch (green) and follow different trajectories in the PCA state space over time. Interestingly, during the late post-stimulus epochs (Figure 3A, darker colors, showing 9 trials with activity in post-stimulus epoch), activity for some trials appears to converge to specific trajectories in the PCA state space. We quantified the number of distinct memory trajectories (over the last second of the post-stimulus period) using k-means. Evaluation of the clustering results was done using the Bayesian Information Criterion (BIC, see Methods, section “Principal component analysis”). We found that for the specific instance (fixed connectivity, see Methods, section “Details of network connectivity”) of the structured network shown in Figure 3, population activity converged to four different robust trajectories (BIC 213.87). These findings suggest that, additionally to the representational shift from the sensory to the memory epoch, a dynamic mechanism is recruited by specific stimuli in an otherwise fixed connectivity network, which allows its rapid re-configuration, in the absence of any learning mechanism, and amounts to the retention of different memoranda.

**Figure 3.**
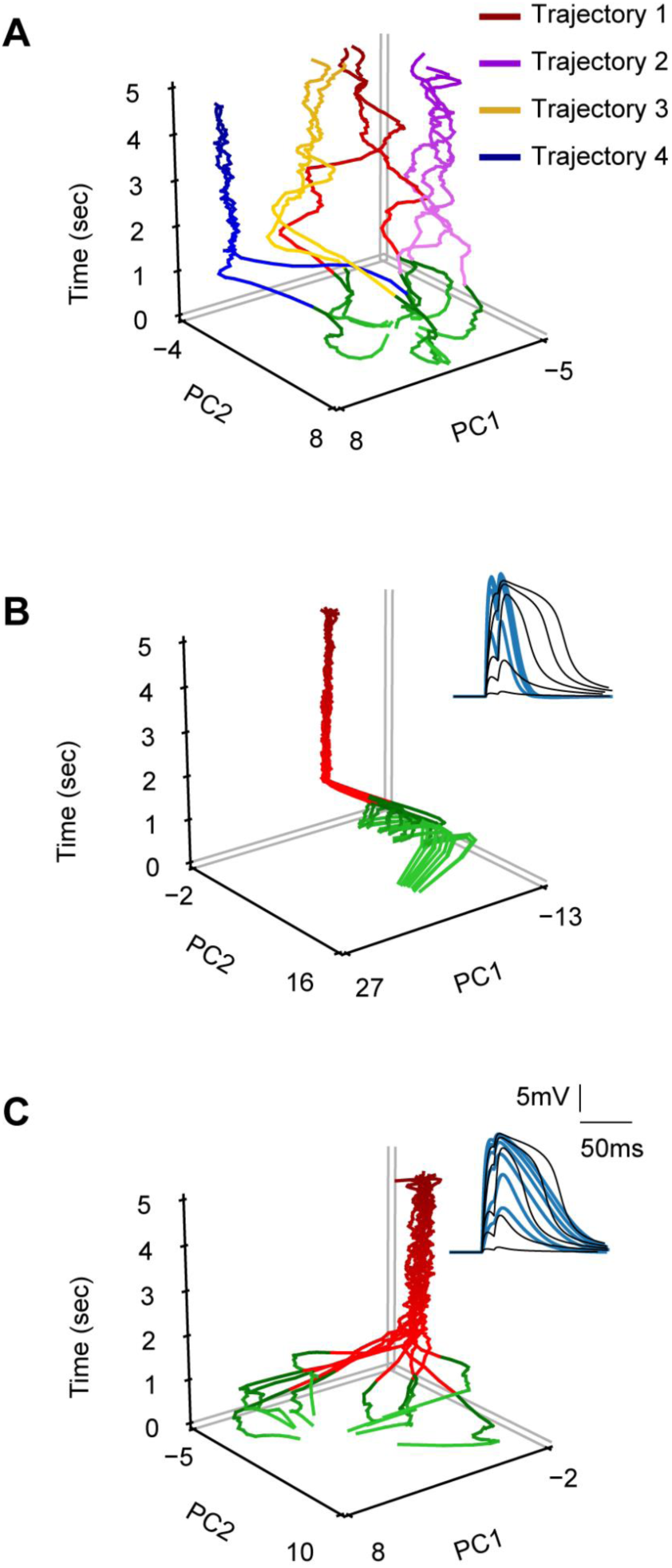
NMDA nonlinearities create distinct stable trajectories. (A) Population activity (n=9 trials with post-stimulus epoch activity) reduced in the first two principal components over time. Stimulus epoch is colored green. Different colors (yellow, red, magenta, blue) correspond to different clusters of post-stimulus activities, as dictated by k-means analysis. Color map luminosity (brighter to darker) denotes time progression. (B) As in A (n=8 trials), with the NMDA conductance set to zero, compensated by increased AMPA conductance. Simulating blockade of NMDARs results in the formation of one trajectory for different stimuli. Inset: voltage response at the soma for the control condition (black) and the NMDA blockade condition (blue). Both the non-linear jump and the depolarizing plateau are eliminated (blue), for increasing synaptic drive when NMDA conductance is zeroed (basal dendrite stimulation, 2 events at 50Hz, activation of 1 to 21 synapses). (C) As in A, after removal of the Mg^2+^ blockage (n=9 trials). This manipulation also results in the formation of a single trajectory. Inset: By removing the Mg^2+^ blockage, the non-linear jump is removed (black control, blue Mg^2+^ blockage), while the depolarizing plateau is retained (basal dendrite stimulation, 2 events at 50Hz, activation of 1 to 21 synapses).

### NMDA nonlinearities underlie multiple memoranda

We hypothesized that a thresholding mechanism acting in short time scales may drive the network activity into the distinct robust trajectories. The most prominent such candidate, in the absence of any short- or long-term plasticity, is the NMDA receptor. To test this hypothesis, we repeated the above simulations while eliminating the NMDA current (conductance set to zero).

To compensate for the reduction in excitability, we increased the AMPA conductance such that the saturated response matched the one when both NMDA and AMPA conductances were included (Figure S2A top, inset in Figure 3B). We further adjusted the network inhibition (see Methods, section “Synaptic manipulations”) to ensure the induction of post-stimulus activity. We found that the AMPA-only model network was unable to express more than one robust trajectory (Figure 3B, BIC=213.2). Activity in the AMPA-only network was also accompanied by a decrease in sparsity (mean of AMPA-only sparsity=0.49) and CV (AMPA-only CV_post-stimulus_= 0.28±0.21, p value=0.0001, comparison with control case, Mann Whitney U test) and an increase in firing frequency (AMPA-only ISI_post-stimulus_=16.6±23.6, p value=0.0001, comparison with control case, Mann Whitney U test) (Figure S2B top, middle). At the population level, there was a reduction in the correlation of the network activity during the stimulus and the post-stimulus epochs as before (Figure S2B bottom, also shown as change in the PCA space, Figure 2B), yet within the stimulus epoch, population activity was stable. These simulations suggest that the NMDA receptor is critical for the expression of multiple robust trajectories and the irregular dynamics in the PFC model network.

Next, we asked which component of the NMDA conductance could be responsible for this finding: its non-linear features or its slow kinetics? The NMDA conductance can create two distinct dendritic phenotypes with increasing synaptic magnitude: a) a non-linear jump in the depolarization, due to the Mg^+2^ mediated non-linearity of the NMDA activation function and b) a depolarization plateau potential following the jump, supported by its slow kinetics (Doron et al., 2017). To assess the contribution of the non-linear jump, we removed the voltage-dependence of the NMDA conductance associated with the Mg^+2^ block, while keeping the slow kinetics and the plateau potential intact (see Figure 3C, Fig S2A bottom). Again, the network failed to support more than one robust trajectory (Figure 3C, BIC=212.9). Note that in this case sparsity is maintained at control levels (sparsity=0.91), while the ISI/CV change significantly (Mg^+2^ block CV_post-stimulus_= 0.55±0.33, p value=0.0001, comparison with control case, Mann Whitney U test, Mg^+2^ block ISI_post-stimulus_= 50.1±91.8, p value=0.0001, comparison with control case, Mann Whitney U test) (Figure S2C top, middle). At the population level, the results were similar to the AMPA-only case (Figure S2C bottom). Manipulations of the kinetics of the NMDA receptor, so as to match those of the AMPA receptor, resulted in identical responses to the AMPA-only simulations (data not shown). Overall, these simulations suggest that while the depolarizing plateau aids in maintaining a sparse network response, it is the non-linear component of the NMDA receptors that serves as a thresholding mechanism for the emergence of multiple activity states.

### Synaptic arrangement along the dendrite regulates the network re-configurability

The spatial arrangement of synaptic inputs is known to affect NMDA receptor opening, resulting in non-linear dendritic integration (Losonczy and Magee, 2006; Poirazi et al., 2003a, 2003b; Polsky et al., 2004). We thus further assessed how synaptic arrangement within the basal dendrites of our model PCs may affect the NMDA-depended generation of multiple trajectories. Specifically, we evaluated how many distinct trajectories can be supported when varying the dendritic locations of recurrent synapses (between PC pairs) and dendritic inhibitory inputs (n=10 different synaptic arrangements). These manipulations resulted in a total of 100 simulated conditions: 10 synaptic arrangements for a given connectivity matrix × 10 trials (note that, for a given trial, when changing the synaptic arrangement, the same PCs are always stimulated). Three examples of the population activity trajectories for different synaptic arrangements, given a specific connectivity matrix and stimulus subpopulation per trial, are shown in Figure 4A-C. In line with our hypothesis, responses in the PCA space vary considerably when changing the synaptic arrangement, giving rise to several distinct activity trajectories. Note that these different trajectories emerge in response to stimulation of the same neurons in a given network (fixed connectivity matrix) with the exact same stimuli. The only difference between trials is the dendritic localization of the incoming inhibitory and recurrent excitatory connections, indicating that dendritic integration is a key player in the observed phenotype.

**Figure 4.**
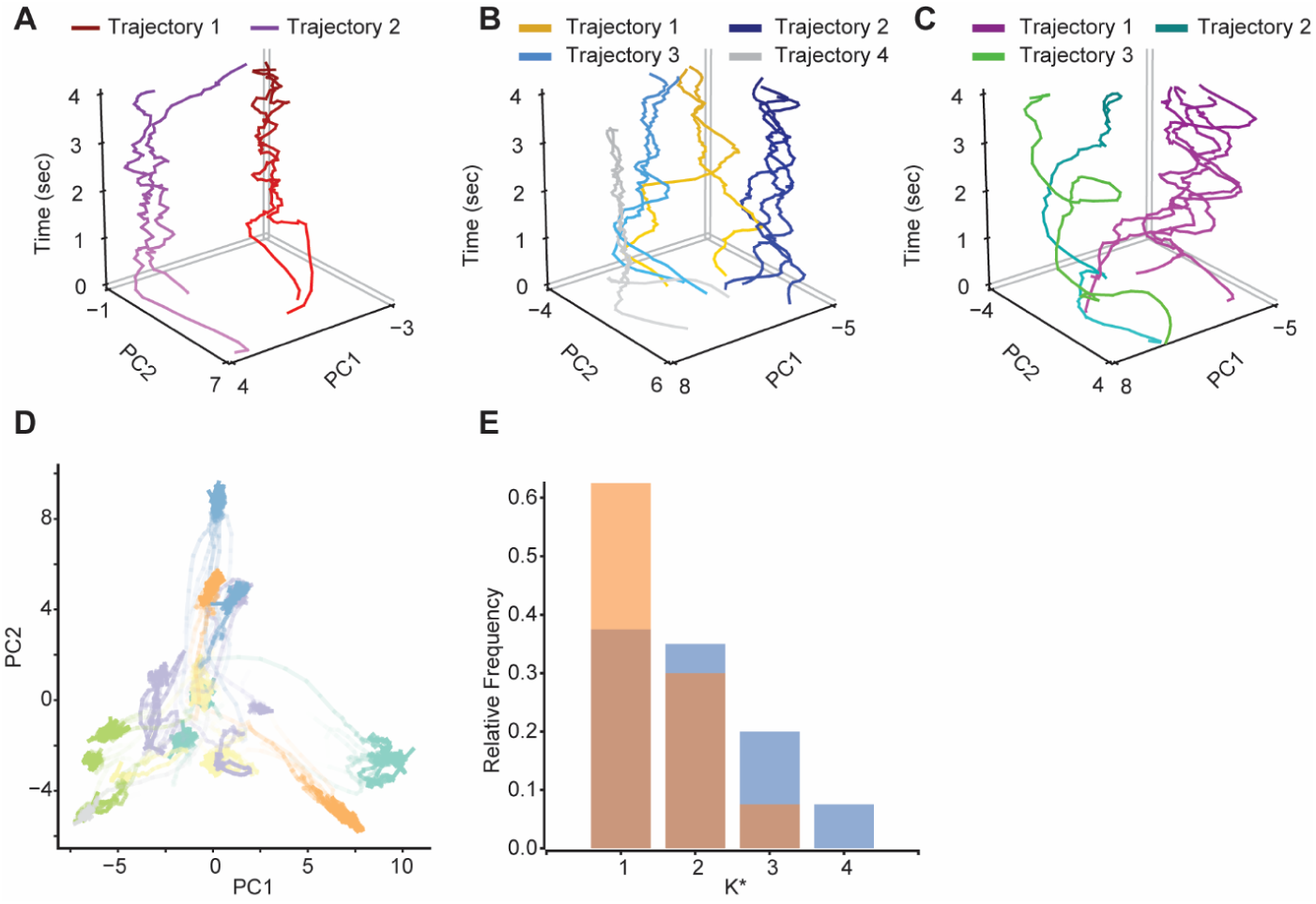
Network activity states in response to same inputs for different synaptic arrangements. (A,B,C) Post-stimulus population activity in PCA space for the same structured network instance, but over three different synaptic arrangements. In the respective trials, the exact same neurons are stimulated. Clusters of trajectories are color coded. It is evident that the number of trajectories changes for each synaptic arrangement. (D) Pooled data for n=10 synaptic arrangements, showcasing that trajectories are mostly nonoverlapping (10 trials, 10 synaptic arrangements). Color code by synaptic arrangement. (E) Histogram of optimal number of clusters (K*) after application of k-means clustering, for all n=4 structured network instances of n=10 synaptic arrangements, for n=10 trials each (blue). In the presence of NMDA nonlinear synaptic integration, a randomly connected network produces fewer trajectories compared to the structured one (orange).

The above findings do not discriminate between the following alternatives: given two synaptic arrangements that support the same number of trajectories, are these identical or distinct in the PCA state space? To answer this question, we pooled all 100 responses (Figure 4D), and found that different synaptic arrangements induce qualitatively different trajectories. These findings suggest that for a given connectivity matrix and input, the arrangement of recurrent synapses enables the network to engage in different subsets of PCA state space, effectively expanding the flexibility of the network.

We investigated the generalization of our results by identifying the number of trajectories found for the 10 different trials, given n=4 different connectivity matrices and n=10 synaptic arrangements (thus amounting to 40 cases). We found that the probability of a network with a given connectivity and synapse arrangement to express more than a single, activity trajectory was 0.625 (sum of probabilities for K* > 1, Figure 4E, blue).

### Structured connectivity exploits NMDA nonlinearities to increase memory

Given the above results, we sought to identify whether structural connectivity can selectively exploit the NMDA non-linearities to induce more than one post-stimulus trajectory. To assess the role of connectivity (both in terms of recurrent connections and the clustering coefficient), we repeated the same simulations on a uniformly connected random network i.e. with each PC pair independently connected with a fixed probability (4 random connectivity matrices, for 10 recurrent synapse arrangements and 10 trials, see Methods, sections “Network Connectivity” and “Synaptic connectivity properties”). Both the structured and the random network types exhibited the same average connection probability (∼0.12), to ensure a fair comparison. In the random case, single-neuron properties were also different compared to the structured one (random ISI_post-stimulus_= 34.7±77.6, p value=0.0001, compared to control case, Mann Whitney U test, random CV_post-stimulus_= 0.56±0.42, p value=0.0001, compared to control case, Mann Whitney U test, Figure S2D top, middle). At the population level, there was a reduction in the correlation of the network activity during the stimulus and the post-stimulus epochs as before (Figure S2D bottom), yet within the stimulus epoch, population activity was more stable compared to the control case. Most importantly, and in the presence of the NMDA conductance, the random network produced fewer distinct trajectories compared to the structured one (Figure 4E). The probability of the random network expressing more than one memory trajectory was 0.375 (sum of probabilities for K* > 1, Figure 4E, orange). These simulations indicate that the structured, highly reciprocal connectivity of the PFC better exploits the NMDA non-linearities to produce additional trajectories compared to a uniform, randomly connected network.

### Distinct trajectories correspond to different, dynamically recruited neuronal assemblies

We showed that the same network model can encode multiple memoranda during the post-stimulus period, utilizing a dynamic thresholding mechanism. How is this mechanism changing the effective connectivity between neurons to create distinct activity patterns? Are these activity patterns associated with specific neuronal assemblies and, if yes, is there a meaningful way that these assemblies are recruited during the post-stimulus epoch? Regarding the first question, we utilized the non-negative matrix factorization (NNMF) approach (Lee and Seung, 1999), which allows us to approximate any neuronal activity pattern over time as a linear combination of a “basis” set. This “basis” set consists of neuronal activity profiles, as opposed to PCA components which do not directly map onto specific neurons (Figure 5A). We applied the NNMF on the network responses for the same network connectivity matrix and synaptic arrangement as used for Figure 3A, for the different trials. Population activity was decomposed to the activities of different neuronal assemblies, whereby the contribution of each assembly has a different weight (i.e. assembly activity) (Figure 5A). Note that assembly recruitment can also be transient during the post-stimulus epoch (Figure 5B), yet each trial can be identified by its assembly composition. Importantly, to assess whether different combinations of assemblies map to different trajectories, we visualized the activity for different trials, color-coded by the trajectory type, where same trajectories appear to be represented by the same assembly combinations (Figure 5C). The engagement of the assemblies in a trajectory can change over time, yet their combination appears distinct between the different trajectories.

**Figure 5.**
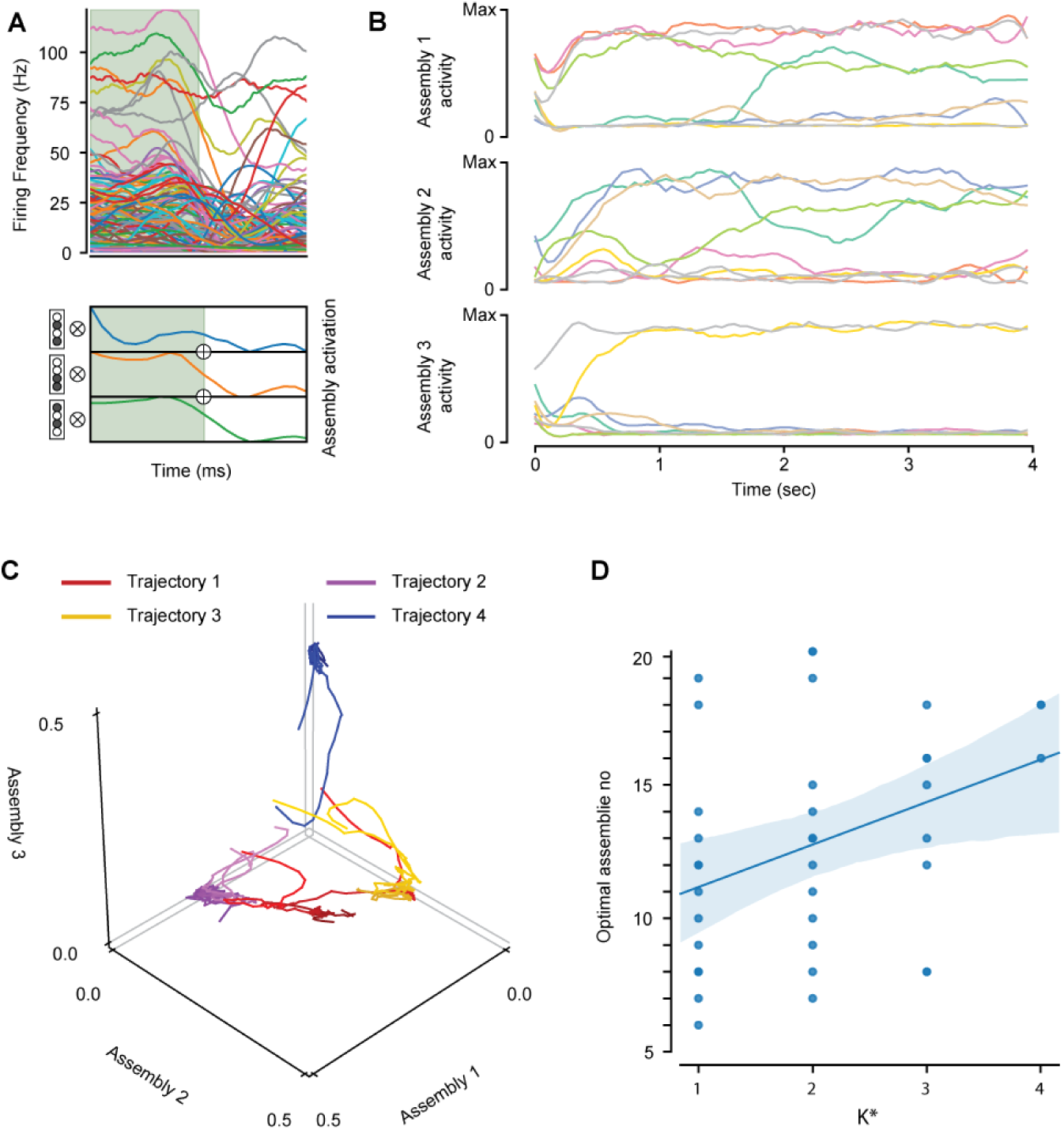
Different neuronal assemblies are active for each population activity trajectory. (A) Top: Smoothed single neuron activities for the last 0.5 sec of stimulus (green area) and the initial 0.5 sec of the post-stimulus epoch of a given trial. Bottom: Same activity, but now reduced with NNMF to 3 components (assemblies). Left: toy examples of assemblies created by 4 neurons. Each assembly is expressed as a vector of participating neurons and each neuron can participate in more than one assembly. Right: each of the assemblies is multiplied (outer product) with its activity vector over time. Population activity is reconstructed by the summation of the respective matrices across assemblies. The dynamic and highly irregular single neuron activity appears smooth in the assembly space and the shift in the representation from the stimulus to post-stimulus epoch is evident. (B) Post-stimulus epoch activity in 9 trials out of 10 with PC activity up until the last second of the simulation, reduced with NNMF to three assemblies and presented each time as the activity of each assembly. Note that assembly activity is dynamic across time. Colors correspond to the different trials. (C) Same data as in B, color coded per the trajectories identified in Figure 3A and plotted in the assembly space. It is evident that trajectories that cluster in PCA space (similar colors), appear to correspond to the activity of specific assemblies/assembly combinations. Note that assemblies are quasi-orthogonal (a neuron can be engaged in more than one assembly) and orthonormal axis are just for visualization purposes. This is expected due to the existence of neurons with mixed selectivity to memoranda. (D) Correlation analysis of optimal number of trajectories and assemblies shows that there is correlation (r=0.37 p=0.017), yet a single trajectory might not necessarily be created from only one assembly.

Concerning the way that these assemblies are recruited during the post-stimulus epoch, correlation analysis of the number of trajectories and the optimal number of assemblies (Methods, section “Non-Negative Matrix Factorization”) for the structured network connectivity, shows interesting results. First, there is a positive correlation (r=0.37, p=0.017) between the optimal number of assemblies and the optimal number of distinct trajectories (Figure 5D). This reveals that trajectories stored in the network tend to be represented by different populations, rather than different firing patterns of the same population. Second, there is no systematic one-to-one correspondence between the number of distinct trajectories and the number of assemblies. This is shown by the identification of more assemblies than trajectories (Figure 5D) and can be interpreted as having multiple assemblies encoding a single trajectory. The latter suggests the presence of assemblies that are reusable across different memoranda and therefore, consisted of neurons that display mixed selectivity. Summarizing, the above results indicate that a recurrent network with NMDA nonlinearities and a structured, highly reciprocal connectivity provide the conditions to have mixed selectivity neurons, instead of the typical randomly connected network. This mixed selectivity is ideal for linearly separating these representations down-stream in complex cognitive tasks, because it stipulates that memoranda representations in network state space are not linearly dependent (can consist of different neurons), thus can more easily be separated by a linear readout neuron (Lindsay et al., 2017; Rigotti et al., 2013). Moreover, these results generate a testable prediction that neurons in PFC, after a task acquisition through learning, form assemblies with mixed selectivity neurons, that can be rapidly utilized, either solo or in combinations in response to the available stimulus and task at hand.

## DISCUSSION

It is not well understood how information entering the PFC can be utilized for flexible decision making: how does a single network represent and support distinct tasks in a rapid, flexible manner (Mante et al., 2013; Stokes et al., 2013; Warden and Miller, 2010)? Experimental evidence indicates that network connectivity can sustain changes in synaptic efficacies faster than classical plasticity, enabling it to rapidly switch between different activity states, driven by input (Fujisawa et al., 2008; Raposo et al., 2014; Stokes et al., 2013) or internal network representations (Saez et al., 2015). Here, we tested the hypothesis that activity switches are the result of NMDA-dependent nonlinear dendritic integration, which is a candidate complementary mechanism to dynamically reconfigure the network connectivity (Arnsten et al., 2010; Fujisawa et al., 2008; Lisman et al., 1998; Stokes, 2015).

We used a state-of-the-art modeling approach that enabled us to dissect the essential mechanisms that drive these highly dynamic phenomena. We built and validated a reduced, compartmental PFC network model that reproduced a large number of experimental observations related to dynamic memory encoding. More specifically, we showed that single neuron responses in the model are irregular, yet together create a coherent low dimensional trajectory. Our findings are consistent with a new experimental view, whereby individual neurons exhibit persistent activity in a highly dynamic manner, while retaining stability and discriminability of WM (Duncan, 2001; Duncan and Miller, 2002; Lindsay et al., 2017; Meyers et al., 2008; Miller et al., 1996; Murray et al., 2017; Rigotti et al., 2013; Saez et al., 2015; Shafi et al., 2007; Stokes et al., 2013). In addition, network activity shifted from encoding of the stimulus, towards a low energy state that represented the memorandum for the specific task at hand, a feature that -to the best of our knowledge-is replicated for the first time, in a single, biologically constrained (and thus recurrent), biophysical network model.

We predict that the key subcellular mechanism responsible for inter-trial changes in dynamic connectivity is the NMDA receptor. Dependent on the spatial arrangement of incoming inputs, the nonlinear integration properties of NMDARs can rapidly shift synaptic efficacies and enable the network to differentially integrate information for short periods of time (Buzsáki, 2010; Stokes, 2015). This is based on several lines of evidence indicating that NMDA receptors are important for PFC function (Lisman et al., 1998; Papoutsi et al., 2014; Wang, 1999; Wang et al., 2013). Furthermore, maximal exploitation of NMDA non-linearities is achieved by connecting subsets of neurons in a clustered, hyper-reciprocal manner, as evidenced in the PFC anatomy (Wang et al., 2006), and previously hypothesized to be beneficial for memory (Brunel, 2016). Although different synaptic (Fiebig and Lansner, 2017; Mongillo et al., 2008) and network (Bouchacourt and Buschman, 2019; Fujisawa et al., 2008; Goldman, 2009; Lindsay et al., 2017; Lundqvist et al., 2016; Rigotti et al., 2010) mechanisms for flexible encoding in local PFC networks were previously hypothesized they fail to holistically summarize the multitude of PFC findings. The key PFC mechanisms we report can create a variable yet discriminable PFC output resembling stable attractor points in a sub-space domain, avoiding the limitations of a linear attractor model (fine-tuned network structure, limited capacity, no flexibility in synaptic weight changes (Goldman, 2009)). Lastly, we point out these mechanisms as a major player in the formation of multiple, transient, assemblies in the PFC. Although neuronal assemblies are not something newly observed (Buzsáki, 2010; Dejean et al., 2016; Fujisawa and Buzsáki, 2011; Peyrache et al., 2009; Schaub et al., 2015), we present a mechanism for rapid assembly creation based on stimulus, with mixed selectivity as a PFC adaptive coding mechanism. Our study provides a mechanistic explanation as to *how* dendritic integration, mediated by the NMDA receptor, can be exploited by the structured connectivity profile of the PFC to provide memoranda representation, encoded stably in time, while maintaining neural variability.

Finally, experimental observations (Saez et al., 2015; Warden and Miller, 2010) point away from the belief that: single neuron = single variable (Sreenivasan et al., 2014), thus increasing the capacity of the network (Barak et al., 2013b; Fusi et al., 2016). The distributed encoding over many neurons helps scale memoranda in a single network, without inducing completely redundant/overlapped responses. In accordance, different (specialized) neuronal ensembles of similarly ‘mixed/tuned’ single neurons can be utilized, as reusable memory units (neuronal ensemble compositionality). This compositionality offers important time-savings when acquiring new tasks, as the network connectivity does not need to be modified by slow, synaptic plasticity processes. It is sufficient to simply activate a new combination of already learned components, thus rapidly augmenting the computational power of the PFC, a region of great behavioral importance (Wang et al., 2019). Our work contributes towards this direction, showing that such a configurability can indeed be supported by a biophysical network model of the PFC, and that such a clustered configuration is indeed optimal in terms of WM storage, contrasted with a non-specific, random projection between neurons, that compromises the compositionality of the network (Barak et al., 2013b).

This work focuses on modelling the key biophysical features of PFC networks, including the extensive connectivity patterns found in experimental PFC works, the dendritic nonlinearity features of pyramidal neurons and the spatial localization of excitatory/inhibitory inputs. Therefore, we refrained from expanding our exploration in many of the free parameters of the model. Specifically, the network size was chosen as to have a minimum of active pyramidals during the post-stimulus epoch, while maintaining the experimentally reported synaptic properties. Simulating networks of greater size is computationally expensive and in our view, does not offer additional insights to this particular problem. To maintain tractability, we modeled pyramidal neurons with just one basal and two apical (proximal/distal) dendrites, validated to reproduce the key synaptic/dendritic nonlinearities in these structures. This reduction is the simplest form of pyramidal neurons that enabled the expression of dynamic phenomena at the network level. We thus expect that more inputs in different dendritic compartments could only extend the model’s adaptive coding complexity (Poirazi et al., 2003b). Moreover, our model included only one inhibitory interneuron type, modeled as a simplified three-compartment node: the soma/proximal dendrite-targeting fast-spiking interneurons. This simplification is clearly not representative of the complex inhibitory networks found in the PFC (Ascoli et al., 2008; Markram et al., 2004; Povysheva and Zaitsev, 2008; Rao et al., 2000; Wang et al., 2004; Zaitsev et al., 2005; Zhang et al., 2017), which also includes dendrite-targeting interneurons and ignores any dendritic nonlinearities that may also contribute to PFC processing (Tzilivaki et al., 2019). Our reasoning for this choice was to focus on excitatory determinants of PFC flexible encoding. Given our own findings about the role of complex interneurons in associative memory (Tzilivaki et al., 2019), we expect that a carefully constrained network model with more realistic interneuronal types will exhibit augmented adaptive coding. Finally, our model did not incorporate any plasticity mechanisms, in order to dissect the contribution of the NMDA nonlinear effects on depolarization from its effects on synaptic plasticity. We believe that our findings regarding the role of NMDA receptors in rapidly configurating network responses are complementary to the slower, plasticity shaped connectivity changes induced by learning. Overall, we are confident that our model is both biologically relevant and tractable, offering significant advantages in probing PFC function.

## Methods

### DETAILS OF MODEL NEURONS

#### The Pyramidal Neuron Model (PC)

The model of the pyramidal cells was adapted from (Papoutsi et al., 2013) (Model DB accession number: 155057). In brief, it consisted of five compartments: the soma, the axon, the basal dendrite, the proximal apical dendrite and the distal apical dendrite (apical tuft). Membrane capacitance on the dendrites was scaled x2 to account for spine membrane area. The somatic, proximal and distal apical dendritic compartments included Hodgkin–Huxley type transient and persistent Na^+^ currents, voltage-dependent K^+^ currents (delayed-rectifier, A-type and D-type), a fast and slow Ca^2+^ -dependent K^+^ current, a hyperpolarization-activated non-specific cation current (h current), and the N-, R-, L- and T-type voltage-dependent Ca^2+^currents (I_caN_; I_caR_; I_caL_; I_CaT_). The basal dendrite included transient and persistent Na^+^ currents, a delayed K^+^ rectifier current, an A-type K^+^ current, a D-type K^+^ current, a N-type Ca^2+^ current and a h current. The axon included a transient Na^+^ current and a delayed rectifier K^+^ current. Indicative electrophysiological properties of the PC model are shown in Table S1. For detailed description see (Papoutsi et al., 2013, 2014).

#### Fast-Spiking interneuron model (FS)

The FS model was adapted from (Konstantoudaki et al., 2014) (Model DB accession number: 168310). In brief, the FS consisted of a somatic, a dendritic and an axonic compartment. The somatic compartment included mechanisms for the slow A-Type K^+^ current, the N-type high-threshold activated Ca^2+^ current and h current. Indicative electrophysiological properties of the FS model are shown in Table S1. For detailed description see (Konstantoudaki et al., 2014).

#### Synaptic Properties

The AMPA and NMDA currents of PCs generated non-linear NMDA-dependent responses at the basal dendrites as in (Nevian et al., 2007), in response to increasing number of activated synapses (2 events at 50Hz, Figure 1A). The AMPA and NMDA EPSC kinetics of the FSs were constrained against data from the respective L5 PFC interneurons (Wang and Gao, 2009), as per (Konstantoudaki et al., 2014). The GABA_A_ conductance was calculated by exponentially interpolating the reported values in (Kubota et al., 2015) (Figure S3). Finally, we implemented a slow inhibitory synaptic current (GABA_B_), as in (Papoutsi et al., 2014) with x2 the conductance value as the GABA_A_ (Table S2). FSs projected to FSs with a single GABA_A_ synapse of 1.7nS as in (Galarreta and Hestrin, 1999). The respective synaptic parameters are listed in Table S2 and detailed description of the synaptic models have been published in (Konstantoudaki et al., 2014; Papoutsi et al., 2013).

### DETAILS OF NETWORK CONNECTIVITY

#### Network Connectivity

The network model consisted of 250 PCs (∼75%) and 83 FSs (∼25%) (Beaulieu, 1993; Dombrowski et al., 2001). We implemented an experimentally-constrained connectivity by treating the biophysical network as a directed graph and its connectivity as a square binary connectivity matrix **A** ∈ ℝ^(M+F)^, where M is the number of PCs and F is the number of FSs. In this matrix, the rows represented neurons’ outgoing connections (efferents) and the columns their incoming connections (afferents), with PCs preceding FSs (for connectivity matrix index i, i ≤ M corresponds to PCs and M < i ≤ M + F corresponds to FSs). Although this matrix contains the connectivity for the whole network, for brevity we will refer to the first M by M part of **A** as its *excitatory* part, containing the PC-PC connectivity. We implemented two different connectivity configurations between the PCs: one highly non-random (termed ‘structured’) and one ‘random’. In the structured configuration a) PC pair connection probability was distance dependent, b) almost half connections were reciprocals, as reported in the PFC (Wang et al., 2006) and c) the PC pair connection probability was re-adjusted to be local-clustering dependent, based on experimental data (Perin et al., 2013; Song et al., 2005). In the random configuration, each pair was connected independently with a fixed probability. Both types of configurations exhibited the same average probability of connection (P_*average*_ ∼0.12), as described below.

In the structured configuration, to implement the distance dependent connectivity constraints, the somata of PCs were regarded as points randomly scattered in a 180×180×180 μm^3^ cubic space (Figure S1A), resulting in a somatic density of 6×10^4^ neurons/mm^3^ (Schuz and Palm, 1989). Distances between these points were calculated by wrapping the cube dimensions in order to eliminate edge effects (Packer and Yuste, 2011; Perin et al., 2011). Each unordered PC pair {i, j}, could be connected in two ways: either unidirectionally (i → j or j → i) or reciprocally (i ↔ j). These disjoint events followed the experimentally reported probabilities as a function of distance in L5 (Perin et al., 2011). In addition, and to reproduce the hyper-reciprocity reported in the PFC, we implemented a similar probability density function (pdf) for both reciprocal and unidirectional connections. The respective pdfs were described as:

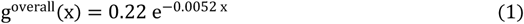

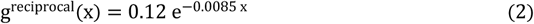

where g^overall^(·) was the pdf of having any connection and g^reciprocal^(·) was the pdf of having a reciprocal connection, both as a function of the distance x (Figure S1B). We derived the unidirectional pdf from their difference:

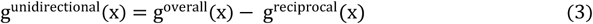

where g^unidirectional^(·) was the pdf of having non reciprocal connection, as a function of the distance x (Figure S1B). Following, for each pair its reciprocal or unidirectional probability of connection, 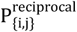 and 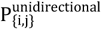 respectively was:

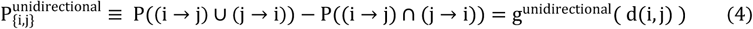

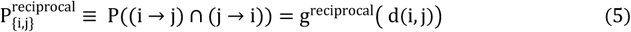

where d(i, j) is the pair’s intersomatic distance x.

The excitatory part of the connectivity matrix **A** was created by rejection sampling for each pair. Specifically, we sampled from a uniform random distribution once for each pair and the excitatory part of the connectivity matrix **A** was determined as:

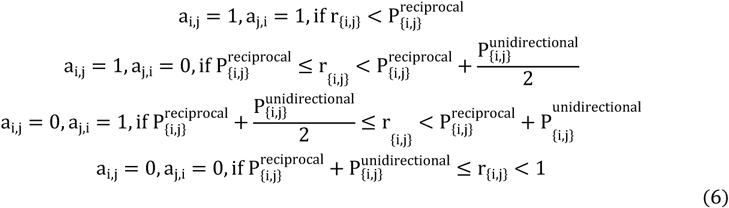

where r_{i,j}_ is a random number on the interval [0,1] and a_i,j_ the elements of **A** matrix. The resulting P_*average*_ was ∼0.12. The P_*average*_ was calculated as the percentage of connected pairs over all pairs,

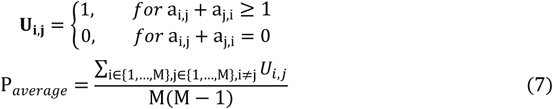

To implement the local-clustering dependent constrains we used the common neighbor rule. According to this rule, pairs sharing increasing number of common neighbors are more probable to be connected (Perin et al., 2011, 2013). A common neighbor is defined as a third PC that receives from or projects to a PC pair. Starting from the above distance dependent excitatory part of connectivity matrix **A**, we iteratively recreated the connectivity to reproduce the common neighbor rule. Specifically, in each iteration we calculated the number of common neighbors for all pairs of the network, resulting in the symmetric matrix **C**_**N**_.

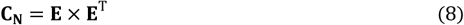

where **E** was the undirected symmetric binary connectivity matrix of the graph of the excitatory part of **A**. Following, we normalized **C**_**N**_

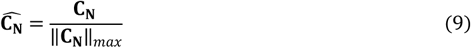

so that the pair of neurons sharing the maximal number of common neighbors is assigned a factor 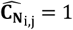 and pairs sharing no common neighbors a factor 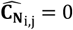 (Perin et al., 2013).

The updated connection probability of a pair 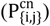 was its 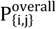, multiplied with 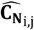

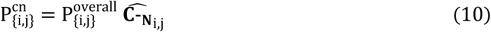

Then, an updated overall connection probability is calculated from the 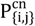 so that the P_*average*_ remains constant:

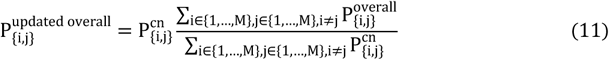

This updated overall connection probability was used to calculate the unidirectional and reciprocal connection probabilities respectively. Towards this, we calculated the distance estimate d′(i, j), that corresponded to the 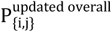 for each pair. The unidirectional and the reciprocal connection probabilities were then calculated using this d′(i, j). The excitatory part of **A**, was recreated with rejection sampling as indicated above.

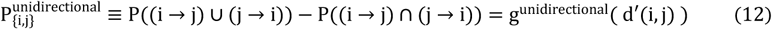

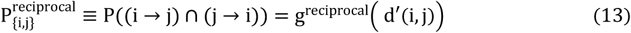

This algorithm ended after 1,000 iterations, where the average clustering coefficient plateaus as in (Perin et al., 2011) (Figure S1C). The clustering coefficient (Watts and Strogatz, 1998) for a single PC was defined as

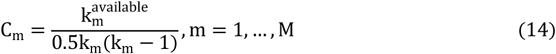

where K_m_ is the number of neighbors (undirected edges to other neurons that neuron M has) and 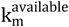 are the undirected edges that exist between these neighbors. The above procedure resulted in the connections shown in Figure S1D,E.

In the case of the randomly connected network, pyramidal neurons were connected with the same independent probability for each pair, irrespective of distance, computed so that the two networks (structured and random) have the same P_*average*_, as defined above. The respective connectivity matrix for the random configuration was created by rejection sampling as indicated above.

We simulated a total of n=4 different network instances (4 structured and 4 random), characterized by their respective connectivity matrix, retaining the interneuron connectivity unchanged (see below). Because this excitatory part of **A** contains only PC connectivity, we changed that without affecting the FS-PC/FS-FS connection probabilities and ratios. These connectivity instances were then translated into the synaptic connectivity of the detailed biophysical models.

For both the structured and the random cases, connectivity between FSs and PCs were distance dependent, as reported experimentally (Packer and Yuste, 2011). Due to the lack of data for PFC regarding the distance-dependent connectivity of L5 FSs with nearby PCs, we used the respective pdf between L5 somatosensory neurons (Packer and Yuste, 2011). Connectivity ratios are summarized in Table S3. FS-to-PC connections were included in **A**_**i**,**j**_, M < i ≤ M + F, j ≤ M and PC-to-FS connections in **A**_**i**,**j**_, i ≤ M, M < j ≤ M + F. Because the overall connection probability reported was conflicting (0.25 in (Otsuka and Kawaguchi, 2009) and 0.67 in (Packer and Yuste, 2011)), we averaged the overall FS-PC connection probability to 0.46. The FS-PC overall, distance dependent connection pdf was:

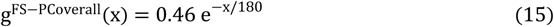

similar to what is reported for somatosensory L5 (Packer and Yuste, 2011). The reciprocal and unidirectional pdfs were based on the FS-PC overall pdf:

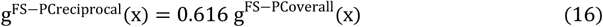

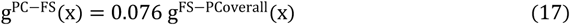

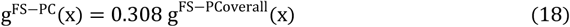

respecting the percentages reported in (Otsuka and Kawaguchi, 2009) (Figure S1F,G,H). The PC-FS connections were added to the connectivity matrix **A** by rejection sampling for each pair, as previously, where the pair {i, j} now refers to FS-PC neurons (M * F pairs in total).

Specifically, we sampled from a uniform random distribution once for each pair and the respective elements of the connectivity matrix, a_i,j_ were determined as:

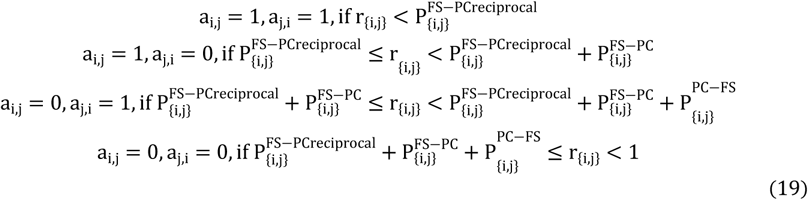

where r_{i,j}_ is a random number on the interval [0,1] and a_i,j_ the elements of **A** matrix (Figure S1F,G,H).

Finally, for the connectivity between FSs-FSs was not distance dependent. We set the connection probabilities 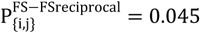 and 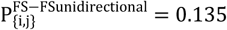, resulting in 0.18 overall connection probability as reported in (Galarreta and Hestrin, 1999) (Table S3). FSs-FSs connections were added to **A** by rejection sampling, where the pair {i, j} now refers to FSs-FSs, with 0.5*F*(*F* − 1) pairs in total:

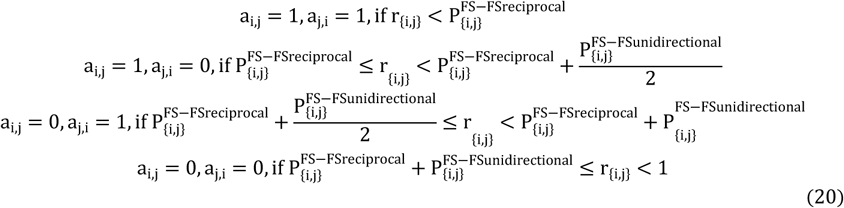

where r_{i,j}_ is a random number on the interval [0,1] and a_i,j_ the elements of **A** matrix. FSs-FSs connectivity elements of **A**, lie on its last F-by-F part with index i, M < i ≤ M + F and j, M < j ≤ M + F.

#### Synaptic connectivity properties

The number and percentage of synapses per type (inhibitory/excitatory) of connection are summarized in Table S4. Single neuron connectivity properties, including the location and the number of synaptic contacts, as well as the latencies between pairs of neurons, were based on anatomical and electrophysiological data. The recurrent connections between PCs were located at the basal dendrite via 5 synapses (Markram et al., 1997) and the synaptic latencies were drawn from a Gaussian distribution with μ = 1.7ms and σ = 0.9ms (Thomson and Lamy, 2007). The precise location of the connecting synapses was random with uniform probability across the basal dendrite. These locations were constant across trials, but reshuffled when the network model underwent a synaptic arrangement. PCs connected to FSs via 2 synaptic contacts (Buhl et al., 1997), with the corresponding latencies drawn from a Gaussian distribution with μ = 0.6ms and σ = 0.2ms (Angulo et al., 1999). FSs project to PCs both to the soma and the basal dendrites (Kubota et al., 2015). Their location was varied randomly with uniform probability between the synaptic arrangements. FSs projected to FSs with a single GABAa synapse as in (Galarreta and Hestrin, 1999) with synaptic latencies drawn from a Gaussian distribution with μ = 1.76ms and σ = 0.07ms (Bacci et al., 2003).

### NETWORK STIMULATION AND MANIPULATION

#### Stimulation

In each trial we stimulated 50 randomly selected (uniform probability) PCs with uncorrelated Poisson inputs (λ=60Hz) for 5sec. The incoming stimulus (40 AMPA and 40 NMDA excitatory synapses), was provided to the proximal apical dendrites of the selected PCs (Kuroda et al., 1998). For the 4 different excitatory parts of the **A** (n=4), 10 synaptic arrangements (defined as randomly positioning the excitatory recurrent PC-PC connections and inhibitory connections along the basal dendrite) were implemented, thus resulting in a total of 40 cases. For each of these cases, we stimulated a total of T=10 times (trial, h), whereby for each trial the 50 stimulated PCs and the stimulus timings were varied. We did not include in the analysis the trials with no activity from any pyramidal neuron in the last second of the post-stimulus epoch.

#### Synaptic manipulations

In order for the network to produce post-stimulus epoch activity we scaled x1.75 the recurrent excitation (g_NMDA_ and g_AMPA_) and x3 the feedback inhibition (g_GABAA_ and g_GABAB_) This was done as the small size of the network limits the number of inputs a single PC received using realistic connectivity probabilities. To compensate for changes in excitability in simulations where g_NMDA_ conductance was set to zero, the g_AMPA_ was further augmented x50. This resulted in the same saturated depolarization when increasing the number of pair-pulse inputs at 50Hz (Figure S2A). Feedback inhibition (g_GABAa_ and g_GABAb_) was scaled to x0.3 so that to allow for post-stimulus activity.

We further manipulated the properties of the NMDA receptors. NMDA current was simulated as in (Polsky et al., 2009):

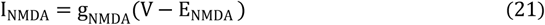

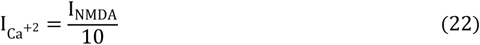

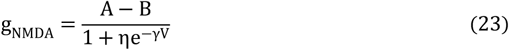

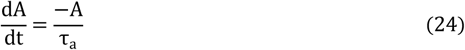

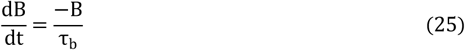

where E_NMDA_ = 0MV, τ_a_ = 90 Ms, τ_b_ = 5 Ms, *Fγ* = 0.08 MV^−1^, η = 0.25MM^−1^.

In order to remove the non-linear ‘jump’ of the NMDA receptors, we removed the Mg^2+^ blockage in (24):

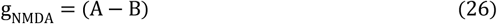

resulting in the saturating depolarization response of Figure S2A (bottom). No other compensations of inhibition were needed in this case to observe post-stimulus epoch activity. Finally, in simulations with the random network connectivity we further adjusted the inhibition to x0.5, in order for the network to produce post-stimulus activity.

### ANALYSIS

#### Network states

We binned PC activity (number of spikes) using discrete temporal windows of length q (q=50ms non-overlapping windows). The instantaneous network state (INS) was defined as the binned activity of each PC in given temporal window, **u** ∈ ℝ^M^. This resulted in 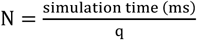 INSs. For a given trial h, we concatenated the N INSs, resulting in a matrix **B**_**h**_ = [**u**_1_, **u**_2_, …, **u**_N_], ∈ ℝ^M×N^, that is the population activity of the trial. We analyzed the population activity across the trials h = 1, …, T, thus resulting in a total of L = TxN INSs. The matrix contained the **B**_**h**_across trials, **D** = [**B**_**1**_, **B**_**2**_, …, **B**_**T**_], ∈ ℝ^M×L^. The latter is referred to as **network state matrix** and characterized the activity of each network instance (same network connectivity and synaptic arrangement, different trials).

#### Cross correlation

Cross correlation between INSs was calculated using the Pearson correlation

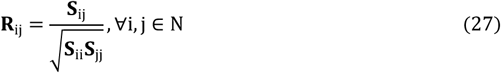

where **S**_ij_ is the covariance of **u**_**i**_with **u**_**j**_.

#### Sparsity

To assess the sparseness of the network response we used the metric proposed by (Rolls, 1997) subtracted from unity, so that higher values correspond to more sparse activity:

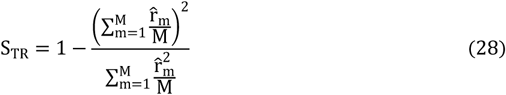

where 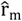 is the average firing rate of the *m-th* PC.

#### Population multidimensional velocity

Network multidimensional velocity **v**_**MD**_ was calculated as the gradient of the activity with respect to time:

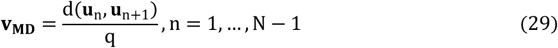

Where d(·) is the Euclidean distance in ℝ^M^ of the two adjacent in time INSs, converted in Hz. The per trial average was used. We further evaluated the change in the average firing frequency of the network using the energy velocity **v**_**E**_, i.e., the change in the overall energy level, across the duration of one trial.

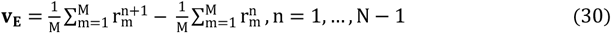

where 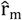 is the firing rate of the *m-th* PC in q.

#### Principal component analysis

In order to examine the neuronal dynamics in a lower dimensional space, the network state matrix **D** was transformed to **F, F** ∈ ℝ^M×L^ using PCA. Following we split the L principal component loadings to the respective trials, representing the *network state space trajectories* per trial. The M_PCA_ principal components with eigenvalues above unity were used for analysis. For visualization, we used the first three principal components.

We evaluated the optimal number of discrete state space trajectories by applying the mean-shift algoritm and k-means, and evaluating the output using the Bayesian Information Criterion (BIC) (Wit et al., 2012). Towards this, from the network state space trajectories (matrix **F**) we kept the last second of each trial 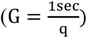 in the matrix 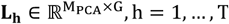. Each trajectory was smoothed prior to clustering with a Savitzky–Golay filter with window size 31 and of polynomial order 3.

We initialized the centroids of k-means using the mean-shift algorithm over all trajectory data and across all trials. We used different Gaussian kernels (K_RBF_) for each trial (thus amounting total to T) characterized by their mean and variance. We used as the mean the average across time of 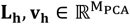.

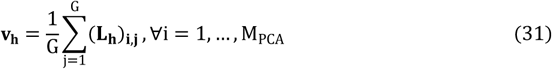

and the variance was set equal to the average variance of **L**_**h**_across all trials. Thus, the K_RBF_ was defined as:

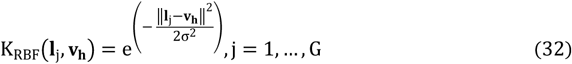

with **I**_**j**_ being the *j-th* column (trajectory point) of **L**_**h**_. We defined the neighborhood of **v**_**h**_ as:

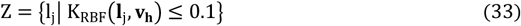

The mean-shift evolves the initial trajectory point **v**_**h**_iteratively and for each iteration the weighted mean was determined as:

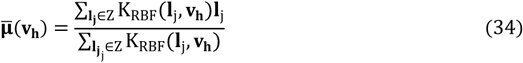

The resulting centroids 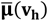 were used as the new mean of the 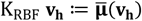 in the next iteration. The iteration continued until centers converge to non-moving positions (translation less than 1/10 of the variance). This process was repeated for all trials h. Following, we assumed K as the optimal cluster number. The T mean-shift centroids were taken in pairs merging those with minimal Euclidean distance, until only K centroids remained (**μ**_**k**_, k = 1, …, K, with 1 ≤ K ≤ T). We finally labeled the average trajectories across time of the trials (**x**_**h**_) to the K clusters. Each average trajectory was assigned to a cluster using the minimum Euclidean distance between **x**_**h**_ and the centroids **μ**_k_, as previously identified by the mean-shift algorithm. Following labels were re-arranged iteratively so that to minimize the intra-cluster Mahalanobis distance:

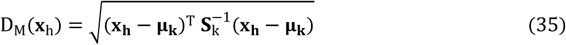

where 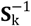 is the inverse of the covariance matrix of the data of cluster k.

Clustering evaluation was done with the BIC:

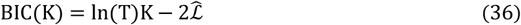

where 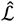 is the value of the maximized log-likelihood function across all T data points (one for each trial), clustered in K clusters:

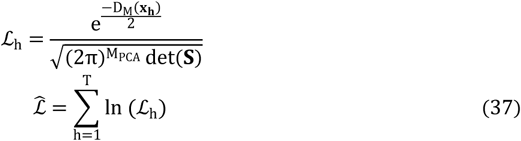

where D_M_(**x**_**h**_) is the Mahalanobis distance of the average trajectory **x**_**h**_ from its cluster centroid and **S** the cluster centroid covariance matrix. We chose the optimal model the one with the smaller BIC value.

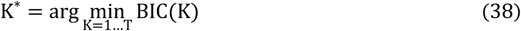

In order to avoid over fitting issues (each trial activity, clustered differently) we set an upper bound on 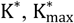 by setting a lower bound on the inter-cluster distance:

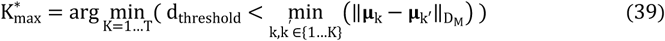

where we set the Mahalanobis distance threshold to d_threshold_ = 3 between cluster centroids **μ**_k_ and **μ**_k′_.

#### Non-Negative Matrix Factorization

To identify the change of effective connectivity between neurons, we used nonnegative matrix factorization (NNMF) as in (Lee and Seung, 1999) on the network instance state matrix **D**. This method utilizes an approximate factorization of the form:

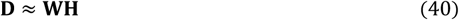

were 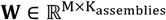 and 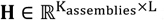 matrices. The matrix **W** corresponds to K_assemblies_ basis patterns (columns of **W**, neuronal assemblies, Figure 5A) and the rows of the matrix **H** contain the coefficients (activity) of each of the assemblies over time.

Since the number of assemblies is dictated by the specific choice of K_assemblies_, we evaluated the optimal number 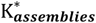 using K-fold Cross-Validation (for a total of 20 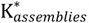). **D** was randomly partitioned into K_CV_ = 20 equally sized subsamples. Of these, a single subsample was retained as the testing data set and the remaining K_CV_ − 1 subsamples were used as the training data set for each K-fold iteration. To partition the matrix, we defined a binary mask **O**, with each of its elements indicating a left out data point 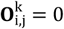 (testing data), or a data point in the training data set 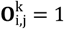, for K-fold iteration k=1,…, K_CV_. This matrix changed for each K-fold Cross-Validation iteration, as predefined by the initial random partition, reflecting the different testing and training datasets. Following, we calculated the **W, H** for each K_assemblies_. We estimated the NNMF factor matrices **W, H** with the multiplicative update rules below as in (Lee and Seung, 2001):

#### Algorithm 1: NNMF with missing elements

1. Initialize W, H randomly
2. **for** t = 1 to 1000 **do**
3. 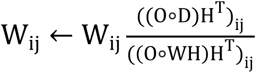
4. 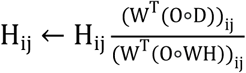
5. **end for**

where ∘ denotes the Hadamard product.

For each of the K-fold iteration, we calculated the error of the model using the testing data set in each K-fold iteration k

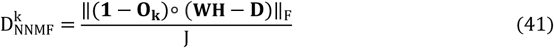

where (**1** − **O**_**k**_) selects the testing data and **W, H** are the NNMF factor matrices of the training data. The error is normalized with J that is the number of zero elements of **O**_**k**_ when calculated for the testing data. Calculating this error for different number of assemblies, creates a ‘bias-variance’ type of error curve, with its minimum indicating the optimal number of assemblies that **D** can be divided to. Due to the random partitioning, the cross-validation procedure was repeated 50 times for the different K_assemblies_. We defined the optimal number of assemblies as the value of 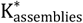 where the average 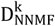 was minimum:

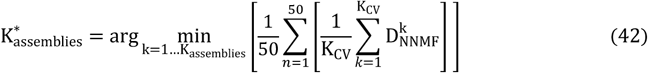

## DATA AND SOFTWARE AVAILABILITY

The microcircuit model was implemented in the NEURON simulation environment (version 7.4) (Hines and Carnevale, 2001). The parallel simulations were executed on a high-performance computing cluster with 312 cores, under CentOS 6.8 environment. Analysis software was written in the Python (version 3.6) programming language. Implementations of algorithms used to compute quantities presented in this study are available at: https://github.com/stamatiad/prefrontal_analysis.

## Acknowledgements

The authors would like to thank Dr. Spyros Chavlis.

## Author Contributions

Conceptualization, S.S.S., P.A., P.P.; Data curation, S.S.S., P.A.; Formal analysis, S.S.S.; Funding acquisition, P.P.; Investigation, S.S.S.; Methodology, S.S.S.; Project administration, P.P.; Resources, P.P.; Software, S.S.S; Supervision, P.A., P.P.; Validation, S.S.S.; Visualization, P.A., P.P.; Writing – original draft, S.S.S, P.A., P.P.

## Declaration of Interests

The authors declare no competing interests.

