## Supplementary Information for "Structured connectivity exploits NMDA non-linearities to enable flexible encoding of multiple memoranda in a PFC circuit model"

**Table S1** Electrophysiological properties of PC and FS models (Related to the Methods section)

| **Cell type** | **Property** | **Value** | **References** |
|---|---|---|---|
| Pyramidal | Input resistance | 80MΩ | (Nasif et al., 2005) |
| Pyramidal | Rheobase | 0.23nA | (Nasif et al., 2005) |
| Pyramidal | AP threshold | -43.5mV | (Nasif et al., 2005) |
| Interneuron | Somatic input resistance | 250.19MΩ | (Zaitsev et al., 2005) |

**Table S2** Synaptic parameters (Related to the Methods Sections)

| **Parameter** | **AMPA** | **NMDA** | **GABA$_A$** | **GABA$_B$** |
|---|---|---|---|---|
| PC, Rise time, ms | 0.6[a] | 5 | 1.5 | 9.8 |
| PC, Fall time, ms | 4.3 [a] | 90 | 14 | 72 |
| FS, Rise time, ms | 0.3 [b] | 0.5 [b] | 3 | - |
| FS, Fall time, ms | 5.5 [b] | 66.3 [b] | 24 | - |
| PC -PC Conductances | 0.19nS [a] | 1.5nS [a] | - | - |
| FS -PC Conductances | - | - | 0.36nS (soma) 0.135nS (average on | 0.72nS (soma) 0.27nS (average on |

| | | | | |
|---|---|---|---|---|
| | | | dendrite) [c] | dendrite) |
| PC-FS Conductances | 0.75nS [b] | 0.32nS [b] | - | - |
| FS-FS Conductances | - | - | 1.7nS [d] | - |

[a] Kinetics fitted to voltage-clamp recordings from (Wang et al., 2008).

[b] As in (Wang and Gao, 2009)

[c] As in (Kubota et al., 2015)

[d] As in (Galarreta and Hestrin, 1999)

[e] To reproduce somatic and dendritic depolarizations as in (Nevian et al., 2007)

**Table S3** Percentage of connected cell pairs (Related to Figure 1).

| **Connection** | **No. of synapses** | **References** |
|---|---|---|
| PC to PC | 12% | (Wang et al., 2006) |
| PC to FS unidirectional | 2% | (Otsuka and Kawaguchi, 2009) |
| PC to FS reciprocals | 15% | (Otsuka and Kawaguchi, 2009) |
| FS to PC unidirectional | 5% | (Otsuka and Kawaguchi, 2009) |

| | | |
|---|---|---|
| FS to FS | 18% | (Galarreta and Hestrin, 1999) |

**Table S4** Number of synapses used in the network (Related to Figure 1)

| **Connection** | **Location** | **No. of synapses** | **References** |
|---|---|---|---|
| Thalamocortical (incoming) | Proximal apical dendrite | 90 | (Kuroda et al., 1998) |
| PC recurrent | Basal dendrite | 5 | (Markram et al., 1997) |
| PC-to-FS | Dendrite | 1 | (Buhl et al., 1997) |
| FS-to-PC | Soma/proximal dendrite | 5 dendrite/3 soma | (Kubota et al., 2015) |
| FS-to-FS | Soma | 1 | (Galarreta and Hestrin, 1999) |

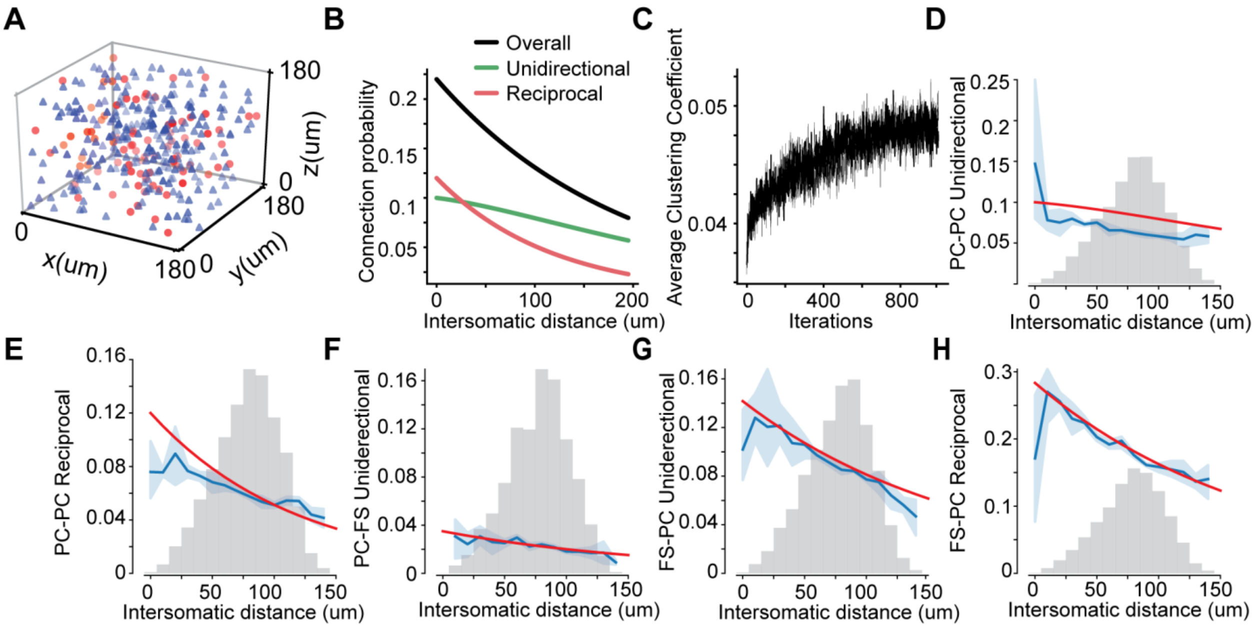


**Figure S1 Structured connectivity (Related to Figure 1 and STAR Methods).**

(A) PCs (blue triangles) and FSs (red circles) were distributed in a 180x180x180 $\mu m^3$ space. (B) Connection probabilities as a function of distance used to create the structured network. (C) Increase of the average clustering coefficient of the structured network, when reforming the connectivity matrix using the common neighbor rule. (D-H) Distance and type dependent connection probabilities of PC and FS neurons in the structured network. Red: probabilities distribution functions based on experimental data (PC-PC adapted from (Perin et al., 2011), PC-FS/ FS-PC/ FS-FS adapted from (Otsuka and Kawaguchi, 2009; Packer and Yuste, 2011)). Blue: relative frequencies of the model network connections per type, across all structured networks instances (n=4, light blue: SE margins across all networks). Light gray: frequency histogram of neuronal pair's distances per type of connection. Due to small number of samples (only a few pairs per bin) SE is greater when moving away from histogram's mean. Probabilities and relative frequencies are displayed on the same axis.

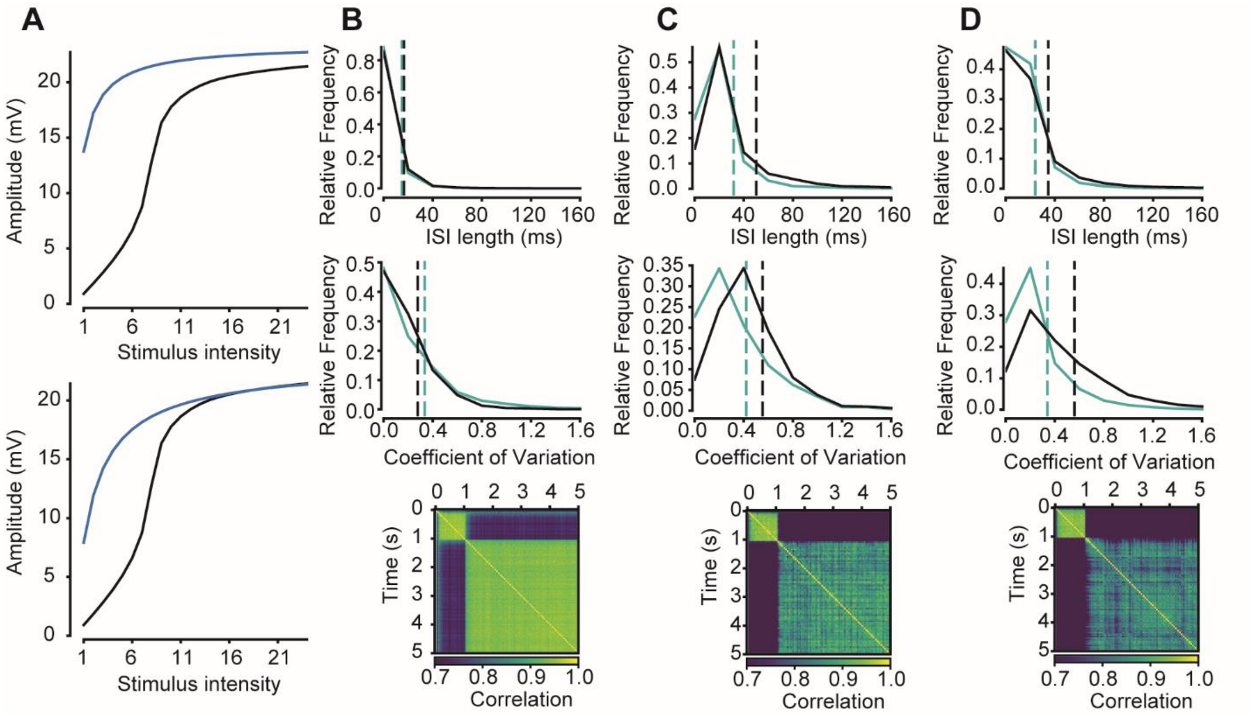


**Figure S2 NMDA non-linearity and its effects on population activity (Related to Figure 3 and STAR Methods).**

(A) Top: Somatic responses to increasing number of synapses (2 events at 50Hz), in the presence of $g_{AMPA}$ and $g_{NMDA}$ (control, black) and in the only AMPA case (blue). The $g_{AMPA}$ in the latter case was adjusted so that the depolarization matched the saturated one of the control case. Bottom: As before, when removing the $Mg^{+2}$ blocking mechanism (blue). (B-D) Top: Stimulus (green) and post-stimulus (black) ISIs (top), CV (middle) and correlation coefficient (bottom), without the $g_{NMDA}$ (B), without $Mg^{2+}$ blockade (C) and (D) for the random network connectivity.

**Figure S3 Related to Figure 3**

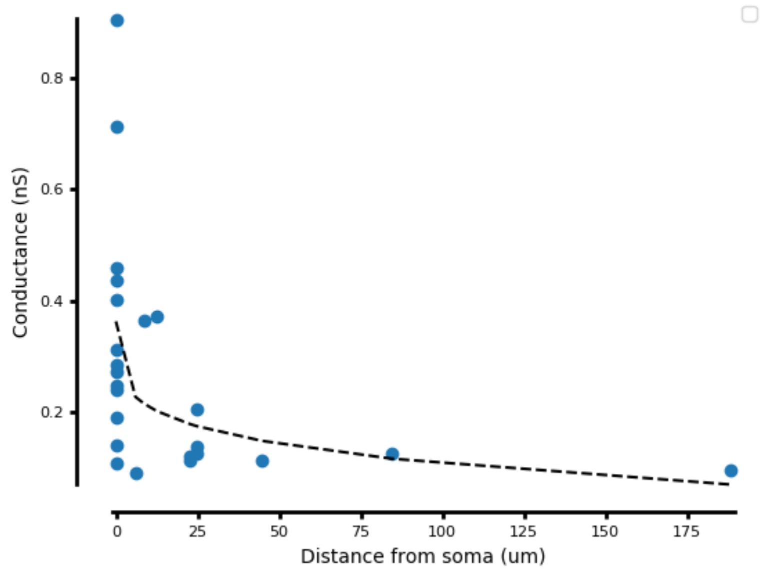


Interneurons projected to pyramidals both in soma and in basal dendrites as reported in (Kubota et al., 2015), with three somatic synapses of 0.36nS (average value) and with 5 dendritic synapses of 0.135nS (average value). The dendritic synapse location varied randomly between synaptic arragements and their conductance calculated by exponentially interpolating the reported values in (Kubota et al., 2015).
